# Aquaporin 1 confers apoptosis resistance in pulmonary arterial smooth muscle cells from the SU5416 hypoxia rat model

**DOI:** 10.1101/2023.10.05.561143

**Authors:** Xin Yun, Shannon Niedermeyer, Manuella Ribas Andrade, Haiyang Jiang, Karthik Suresh, Todd Kolb, Mahendra Damarla, Larissa A. Shimoda

## Abstract

Pulmonary arterial hypertension (PAH) is a deadly condition that arises from increased pulmonary vascular resistance due to contraction and remodeling of the pulmonary arteries. The structural changes that occur in the pulmonary arteries include thickening of the medial (smooth muscle) layer resulting from increased proliferation and resistance to apoptosis. The mechanisms underlying apoptosis resistance in PAH are not fully understood. In cancer cells, high expression of aquaporin 1 (AQP1), a water channel, is associated with apoptosis resistance. We previously showed functional AQP1 protein was expressed in pulmonary arterial smooth muscle cells (PASMCs) and was upregulated in pre-clinical models of pulmonary hypertension. Whether AQP1 controls susceptibility of PASMCs to apoptosis in pre-clinical models of PAH is unknown. In this study, we used PASMCs isolated from control rats and rats exposed to SU5416 plus hypoxia (SuHx) to test the role of AQP1 in modulating apoptosis in PASMCs. We found that elevated levels of AQP1 in PASMCs from pulmonary hypertensive rats were necessary for resistance to apoptosis, and that apoptosis resistance could be conferred by increasing expression of AQP1 in PASMCs from control rats. Moreover, in exploring the downstream pathways involved, we found AQP1 levels influence the expression of Bcl-2, with enhanced AQP1 levels corresponding to increased Bcl-2 expression, resulting in reductions in the ratio of BAX to Bcl-2 as are typically associated with apoptosis resistance. These early results provide a mechanism by which AQP1 can regulate PASMC fate and suggest further investigation could provide additional clues regarding whether AQP1-mediated apoptosis resistance contributes to PAH development or progression and whether AQP1 might be a suitable target for therapy.

## Introduction

Pulmonary arterial hypertension (PAH) is a life-threatening condition that can arise from various etiologies. Regardless of the inciting cause, increased pulmonary vascular resistance due to contraction and remodeling of the pulmonary arteries leads to increased afterload on the right ventricle, ultimately resulting in right ventricular failure. Current approved therapies provide symptom relief and can slow disease progression by inducing vasodilation, but none specifically target the structural remodeling that is a key feature of PAH. Thus, there are no curative therapeutic options.

The vascular remodeling observed in both patients with PAH and animal models includes intimal and medial thickening due to cell proliferation, extension of muscle into non-muscular pre-capillary arterioles, likely due to migration of PASMCs and endothelial-to-mesenchymal transdifferentiation^1-4^. While these processes are prevalent in the early stages of disease and contribute to vascular narrowing, it is increasingly recognized that as the condition progresses, resistance to apoptosis and failure of cell turnover becomes a significant contributor^5-7^. Normally, apoptosis is a highly regulated process of cell death that maintains homeostasis by orderly removal of non-functioning or damaged cells. Impaired apoptosis, as observed in many types of tumor cells^8^,^9^, leads to uncontrolled proliferation and tumor growth. In the case of PAH, resistance to apoptosis has been reported in PASMCs from patients^5 6,10^ and is believed to contribute to vascular narrowing and/or complete obliteration of the arterial lumen.

The mechanisms underlying apoptosis resistance in PAH are not fully understood. In cancer cells, several reports linked excess expression of the protein, aquaporin 1 (AQP1), with apoptosis resistance^11-15^. AQP1 is a water channel that is expressed in a variety of cell types and tissues, including red blood cells, endothelium, renal tubules, tumor cells and sweat glands^16^. We showed functional AQP1 protein was expressed in PASMCs^17^, and was upregulated in pre-clinical models of pulmonary hypertension (PH), including the chronic hypoxia^17^ and SU5416 plus hypoxia (SuHx)^18^ rat models. Loss of AQP1 attenuated the development of PH in response to chronic hypoxia^19^, and we showed AQP1 was required for PASMC migration and proliferation via a mechanism involving the C-terminal tail and upregulation of β-catenin^20, 21^. Whether AQP1 controls susceptibility of PASMCs to apoptosis in the SuHx model is unknown, although in unstimulated control human PASMCs^22^ and in murine PASMCs subjected to in vitro hypoxia^19^, loss of AQP1 increased caspase 3/7 activity, which could be consistent with increasing apoptosis.

The mechanism by which AQP1 might regulate apoptosis is unclear. When cells are subjected to stress, via mitochondrial dysfunction, oxidative damage, stimulated proliferation or exposure to apoptotic stimuli, a series of well characterized events are set into motion, including suppression of Bcl-2, an inhibitor of the pro-apoptotic molecule, BAX^23^. Upon activation, BAX homo-oligomerizes and translocates to, and permeabilizes, the mitochondrial membrane, allowing release of cytochrome C (cyt C) from damaged mitochondria^24^. Cytosolic cyt C then activates caspase 3, resulting in DNA fragmentation and condensation, cell blebbing and formation of apoptotic bodies disposed of through phagocytosis. In cancer cells, AQP1 has been proposed to be upstream of the Bcl-2 pathway^25^,^26^. Whether Bcl-2 and BAX expression are regulated by AQP1 in PASMCs is unknown.

In this study, we tested the hypothesis that enhanced AQP1 expression in PASMCs from the well-established SuHx rat model of PAH contributes to apoptosis resistance due to reciprocal regulation of BAX and Bcl-2, with high AQP1 levels causing an increase in Bcl-2 and repression of BAX. We tested this hypothesis using gain and loss of function experiments in PASMCs isolated from normoxic control rats and from the SuHx model.

## Methods

All protocols were reviewed by and performed in accordance with Johns Hopkins University Animal Care and Use Committee. Protocols and procedures comply with NIH and Johns Hopkins Guidelines for the care and use of laboratory animals.

### Sugen/Hypoxia model

Pulmonary hypertension was induced in adult male Wistar rats (150 –200 g; Harlan Farms) by a single subcutaneous injection of SU5416 (20 mg/kg), prepared in a carboxymethylcellulose (CMC)-containing diluent as described previously^27^, followed by exposure to 10% O_2_ (hypoxia) for 3 wk. The rats were exposed to normoxic conditions for <5 min twice a week to change cages and replenish food and water. At the end of 3 wk, rats were returned to normoxia for an additional 2 wk. Control rats were injected with vehicle and maintained in room air (normoxia) for 5 wk. All animals were kept in the same room and exposed to the same light-dark cycle and ambient temperature and were housed in standard rat cages (3 rats/cage) with free access to food and water. At the end of exposures, rats were anesthetized (Ketamine, 75 mg/kg; Xylazine, 7.5 mg/kg) and depth of anesthesia was confirmed via paw pinch prior to measurement of right ventricular systolic pressure (RVSP) via transdiaphragmatic right heart puncture with a heparinized 23 gauge needle attached to a pressure transducer for 1-3 min. RVSP for each animal was measured as the average of at least 5 continuous heartbeats in animals with a minimum heart rate of >200 bpm. Exposure to the SuHx protocol resulted in an increase in RVSP (**Fig 1A**). Following measurement of RVSP, animals were euthanized via exsanguination and the heart and lungs removed and transferred to a dissecting dish filled with cold N-[2-hydroxyethyl]piperazine-N’-[2-ethanesulfonic acid] HEPES-buffered salt solution (HBSS) containing (in mmol/L): 130 NaCl, 5 KCl, 1.2 MgCl_2_, 1.5 CaCl_2_, 10 HEPES, and 10 glucose, with pH adjusted to 7.2 with 5 mol/L NaOH. Under a microscope, the atria and large conduit vessels were removed and the right ventricle (RV) wall was carefully separated from the left ventricle and the septum (LV+S). Both portions were blotted dry and weighed. As shown in **Fig 1B**, RV/LV+S weight ratio was greater in SuHx rats compared to normoxic rats. Coupled with the increase in RVSP in SuHx rats, these results indicate development of pulmonary hypertension in our model.

**Figure 1:**
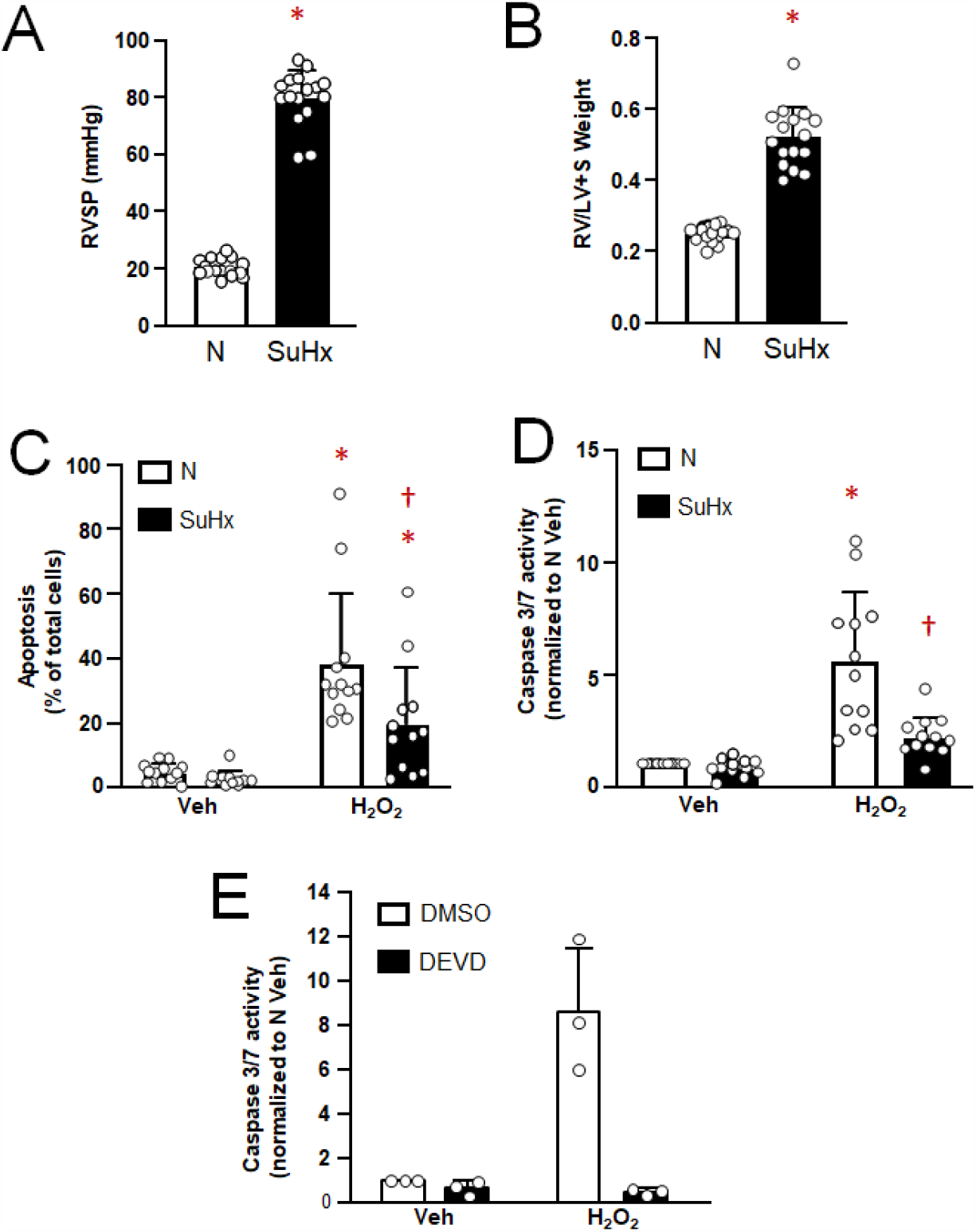
Apoptosis in pulmonary arterial smooth muscle cells (PASMCs) from normoxic (N) and Su5416+hypoxia (SuHx) rats. **A-B)** Bar and scatter graphs represent mean ± SD values and individual values (n=16 rats for each group) for **A)** right ventricular systolic pressure (RVSP) and **B)** right ventricle to left ventricle plus septum (RV/LV+S) weight in N and SuHx rats. * indicates significant difference (p<0.05) from N value via t-test. **C and D)** Bar and scatter graphs represent mean ± SD values and individual values (n=12 rats per group) for apoptosis measured via **C)** Hoescht staining and D) caspase 3/7 activity assay in PASMCs from N and SuHx rats challenged with vehicle (Veh; PBS) or H2O2 (500 μM; 24 h). Values are presented as percent of total cells. For **C** and **D**, * indicates significant difference (p<0.05) from Veh value in the same condition; † indicates significant difference (p<0.05) from H_2_O_2_ value in cells from N rats Significance assessed by 2-way ANOVAwith Holm-Sidak post-hoc test. **E)** Bar and scatter graphs represent mean ± SD values and individual values (n=3 per group) for caspase 3/7 activity in PASMCs from N rats challenged with Veh or H_2_O_2_ in the presence of DMSO (vehicle) or the caspase 3/7 inhibitor DEVD (50 mg/ml)

### Isolation and culture of PASMCs

Resistance level pulmonary arteries (200-600 μM outer diameter) were isolated, the adventitia removed and arteries opened for endothelial denudation with a cotton swab. Tissue was placed in 4°C HBSS for at least 30 min, transferred to room temperature reduced-Ca^2+^ HBSS (20 μm CaCl_2_) for at least 20 min and then digested in reduced-Ca^2+^ HBSS containing: type I collagenase (1750 U/ml), papain (9.5 U/ml), bovine serum albumin (2 mg/ml), and DTT (1 mM) at 37°C for 15-20 min. After digestion, the tissue was transferred to Ca^2+^-free HBSS in a small round bottom tube and slowly pipetted up and down to create a single cell suspension. Dispersed PASMCs were cultured in Smooth Muscle Cell Medium (Sciencell, Cat 1101) supplemented with smooth muscle cell bullet kit (Lonza) with 1% penicillin-streptomycin to expand cell numbers. All cells were used at passage 1-2 and were growth arrested in smooth muscle basal media (Lonza) supplemented with 0.3% FBS for 24-48 h before beginning experiments. Assessment of smooth muscle culture purity was performed using: 1) [Ca^2+^]_i_ responses to 80 mM KCl and 2) immunofluorescence in cells stained with smooth muscle-specific α-actin (SMA; 1:400, A2547, Sigma) and calponin (1:100, ab700, Abcam) or smooth muscle myosin heavy chain (SMMHC; 1:100, ab125884, Abcam) and DAPI (nuclear stain; 1:10,000 in PBS, Invitrogen). Only cultures where >90% of cells exhibited at least 50 nM increase in [Ca^2+^]_i_ and/or positive SMA and calponin or SMMHC expression were used for these studies.

### Hydrogen peroxide (H_2_O_2_) exposures

To induce apoptosis, cells were exposed to H_2_O_2_ (Sigma). At approximately 60% confluence, cells were placed into basal media for 24 h. Cells were then washed 3 times with PBS before being placed in serum-free media containing freshly made H_2_O_2_ (500 μM) and incubated for 24 h before additional measurements (apoptosis, caspase activity or immunoblot) were made.

### Hoechst staining

PASMCs were stained with Hoechst 33342 dye (3 μL/ml of 1:100 dilution; 62249; Invitrogen) for 15-30 min at 37° C to visualize apoptotic cells. Apoptosis was identified by observing alterations in chromatin morphology (i.e., condensation) upon staining. For each sample, 9 images were captured at 20X via fluorescent microscopy and analyzed using ImageJ. Apoptosis was determined as the percent of apoptotic cells over total cells and counts for each image within a sample were averaged to obtain a single value.

### Caspase activity assay

Caspase 3/7 activity was measured using the Caspase-Glo® 3/7 Assay System (G8090, Promega) per the manufacturer’s instructions. Briefly, equal amounts of cell lysate (2.5 or 5 μg) and T-PER (ThermoFisher) for each sample were placed into the wells of a 96 well plate in duplicate. Proluminescent caspase-3/7 DEVD-aminoluciferin substrate was added to each well and read with a luminometer at 5 min intervals up to 60 min. Luminescence values at within the linear portion of the curve (typically at 30 min) were background subtracted and normalized to control.

### Western Blotting

PASMCs were washed with PBS, and total protein extracted in ice-cold T-PER buffer containing protease inhibitors (Roche Diagnostics). Proteins were quantified by use of the BCA protein assay (Pierce) and 5 μg total protein was resolved by 10% SDS-PAGE gels transferred onto polyvinylidene difluoride membranes, which were then blocked with 5% nonfat dry milk in Tris-buffered saline containing 0.2% Tween 20. Membranes were probed with primary antibodies AQP1 at 1:4,000 (AQP11A, Alpha Diagnostic Intl. Inc.), Bcl-2 at 1:1,000 (ab196495, abcam) or BAX at 1:1000 (ab32503, abcam). Antibodies are verified with siRNA knockout by our lab previously^17^ or the antibody company. Bound antibodies were probed with horseradish peroxidase-conjugated anti-rabbit or anti-mouse IgG (1:10,000, 52200336 and 52200341, Kirkegaard & Perry Laboratories) and detected by enhanced chemiluminescence using a Chemidoc (BioRad) gel imaging system. Membranes were then stripped and re-probed for β-tubulin (1:10,000, T7816, Sigma) as a housekeeping protein. Protein levels were quantified by densitometry using ImageJ.

### Adenovirus Infection

Adenoviral constructs containing a hemagglutinin (HA)-tagged wild-type AQP1 (AdAQP1) and GFP were created as described previously^20^. PASMCs were placed in basal media and infected with virus (50 ifu/cell) for 48 hr before being used in protein expression measurements and functional assays. Cells infected with the same adenovirus containing GFP (50 ifu/cell) were used as controls. We previously demonstrated expression and appropriate localization of AQP1 expressed using this adenovirus^20^.

### siRNA

Depletion of endogenous AQP1 was achieved using siRNA specifically targeting AQP1 (siAQP1) and non-targeting (siNT; control) siRNA obtained as a “smart pool” (Horizon Discovery) as previously described^17^. PASMCs were incubated with 100 nM of siRNA for 16 h in serum- and antibiotic-free media, after which serum was added to media for a total concentration of 0.3% FBS. Cells were incubated under these conditions for 24 hr, and then media was replaced and cells were incubated for an additional 48 hr in basal media (0.3% FBS) prior to experiments.

### Statistical analysis

Data are expressed as scatter plots with bars representing means ± SD. Each dot represents a separate experimental run, and since all experimental runs were performed on tissue/cells from different animals, “n” also refers to the number of animals. All data were tested for normality and equal variance prior to running statistical tests. Data that were not normally distributed were log(10) transformed and retested for normality and equal variance prior to running statistics. Statistical comparisons were performed using Students t-test for data in two groups, or one- or two-way ANOVA with a Holm-Sidak post hoc test for multiple group comparisons.

## Results

### Effect of hydrogen peroxide on PASMCs

In cells isolated from normoxic rats, basal rates of apoptosis, measured via Hoescht staining, averaged less than 5% (**Fig 1C**). The average rate of basal apoptosis in PASMCs from SuHx rats was lower than, but not statistically different from, that measured in PASMC from normoxic rats. With application of the apoptotic stimulus, H_2_O_2_ (500 μM) for 24 hr, the rate of apoptosis increased in normoxic cells to approximately 40%. In PASMCs from SuHx rats, the increase in apoptosis induced by H_2_O_2_ was significantly less (approximately 20%) than that observed in normoxic cells, indicating that SuHx cells have reduced susceptibility to apoptotic stimuli.

To complement results obtained using Hoescht staining, we also performed caspase 3/7 activity assays. Consistent with our results with Hoechst staining, basal caspase 3/7 activity in unstimulated cells was not different between PASMCs from normoxic and SuHx rats (**Fig 1D**). As expected, challenge with H_2_O_2_ (500 μM) for 24 hr resulted in a marked increase in caspase 3/7 activity in normoxic cells. In SuHx cells, caspase 3/7 activity after exposure to H_2_O_2_ was significantly lower than that observed in H_2_O_2_-exposed normoxic cells and not significantly different from SuHx PASMCs under unstimulated conditions.

To verify the caspase 3/7 activity measures using the CaspGlo assay truly reflected caspase 3/7 activity, we tested the effect of the specific inhibitor of caspase 3/7, DEVD. We repeated H_2_O_2_ exposure in cells from normoxic animals in the presence of DEVD (50 mg/mL) or vehicle (DMSO) (**Fig 1E**). The addition of DEVD significantly reduced caspase 3/7 activity, indicating the luminescence measured was indeed due to activity of caspase 3/7.

### Role of increased AQP1 protein in apoptosis susceptibility

We found that PASMCs from our SuHx rats exhibited increased AQP1 protein levels compared to controls (**Fig 2A**), consistent with our previously published results^18^. Given the inverse correlation between AQP1 protein abundance and apoptosis reported in several tumor cell types^12, 28-30^, we tested whether elevated levels of AQP1 could contribute to apoptosis resistance in SuHx PASMCs. Compared to cells treated with non-targeting siRNA (siNT), knockdown of AQP1 using siRNA (siAQP1) significantly reduced, but did not fully deplete, AQP1 protein in SuHx cells (**Fig 2B**). With knockdown of AQP1, the percent of cells from SuHx rats undergoing apoptosis at baseline (vehicle-treated cells) was greater than in cells treated with siNT, although the difference did not reach statistical significance (p=0.3431; **Fig 2C**). In response to H_2_O_2_, SuHx PASMCs treated with siNT were resistant to apoptosis, with a small increase in the percent of apoptotic cells that did not reach statistical significance compared to vehicle treatment. In contrast, depletion of AQP1 restored the ability of H_2_O_2_ to induce apoptosis in SuHx cells (**Fig 2C**). As shown in Fig 1, H_2_O_2_ has little effect on caspase 3/7 activity in SuHx cells. Consistent with our apoptosis results, caspase 3/7 activity in response to H_2_O_2_ was also significantly increased when AQP1 was depleted in SuHx cells (**Fig 2D**.

**Figure 2:**
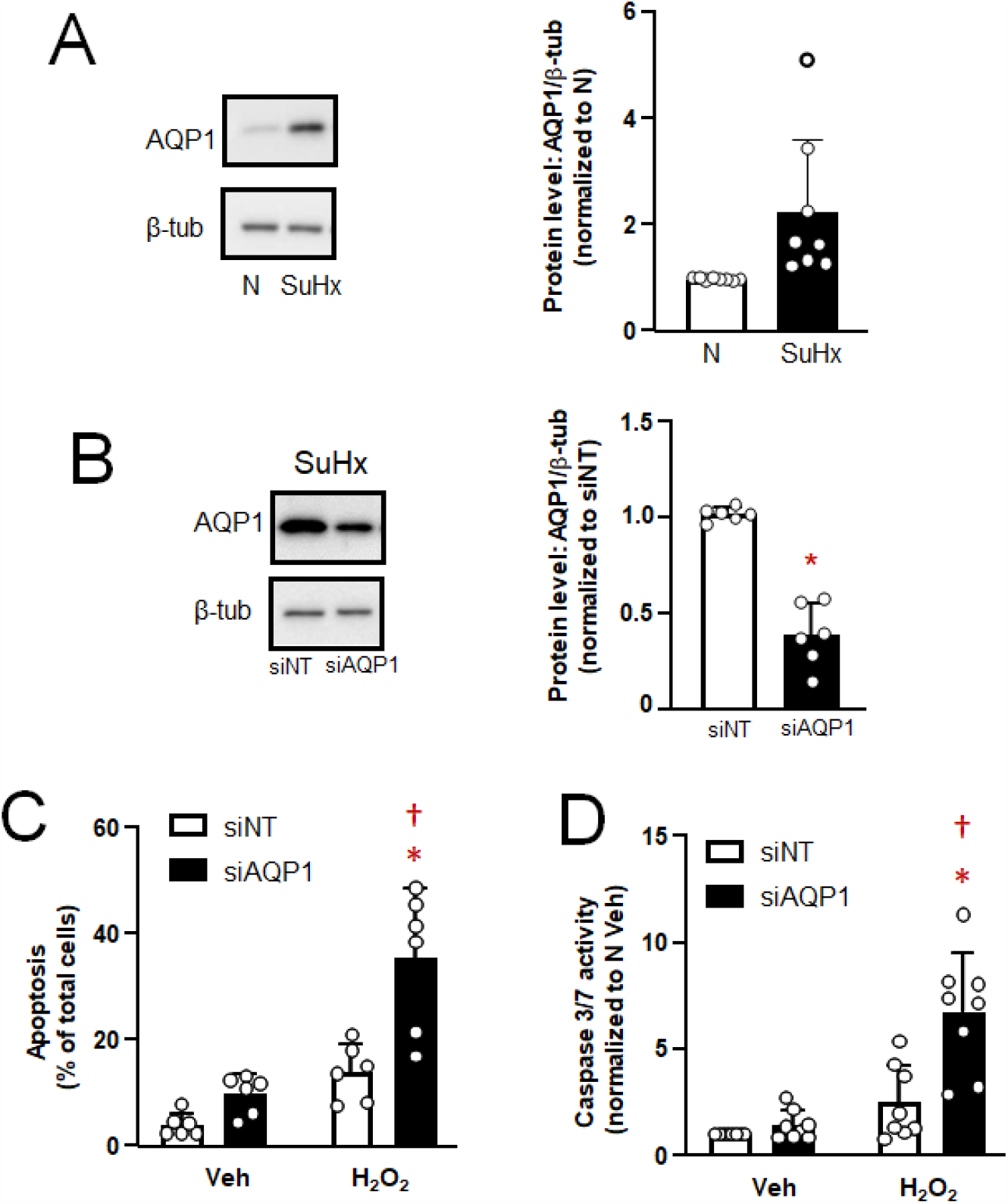
Effect of AQP1 depletion on apoptosis in pulmonary arterial smooth muscle cells (PASMCs). **A)** Representative blots show AQP1 protein levels in PASMCs from normoxic (N) and Su5416/hypoxia (SuHx) rats. Bar and scatter plot graphs represent mean ± SD values and individual values (n=8 rats per group) for AQP1 protein levels. * indicates significant difference (p<0.05) from N value by t-test. **B)** Representative blots show siRNA targeted to AQP1 (SiAQP1) reduced AQP1 protein levels in PASMCs from SuHx rats compared to cells transfected with a non-targeting siRNA (siNT). Bar and scatter plot graphs represent mean ± SD values and individual values (n=6 rats per group) for AQP1 protein levels. * indicates significant difference (p<0.05) from siNT value by t-test. **C and D)** Bar and scatter graphs represent mean ± SD values and individual values for apoptosis measured via **C)** Hoescht staining (n=6 rats per group) and **D)** caspase 3/7 activity assay (n=8 rats per group) in PASMCs from SuHx rats transfected with siNT or SiAQP1 and challenged with vehicle (Veh; PBS) or H_2_O_2_ (500 μM; 24 h). Values for apoptosis are presented as percent of total cells while values for caspase 3/7 activity are normalized to siNT Veh cells. * indicates significant difference (p<0.05) from siAQPI cells treated with Veh; † indicates significant difference (p<0.05) from H_2_O_2_ value in cells treated with siNT.

### Effect of increasing AQP1 expression on apoptosis susceptibility in control PASMCs

Since reducing AQP1 levels in SuHx restored susceptibility to an apoptotic stimulus, we next tested whether increasing AQP1 levels in control cells could confer resistance to apoptosis. Using an adenovirus encoding wild-type AQP1, we were able to increase AQP1 protein levels in control PASMCs (**Fig 3A**). In these experiments, a lower band was visible in all lanes, corresponding to native AQP1. In samples infected with AdAQP1, a second band of slightly higher weight was clearly visible, corresponding to the expressed HA-tagged AQP1. Increasing AQP1 protein levels in control cells had no significant effect on percent of cells undergoing apoptosis (**Fig 3B**) or caspase 3/7 activity (**Fig 3C**) at baseline. Stimulation with H_2_O_2_ increased both apoptosis (**Fig 3B**) and caspase 3/7 activity (**Fig 3C**) in cells infected with AdGFP, as expected, whereas increasing AQP1 levels with AdAQP1 suppressed but did not completely prevent H_2_O_2_ from inducing apoptosis or increasing caspase 3/7 activity.

**Figure 3:**
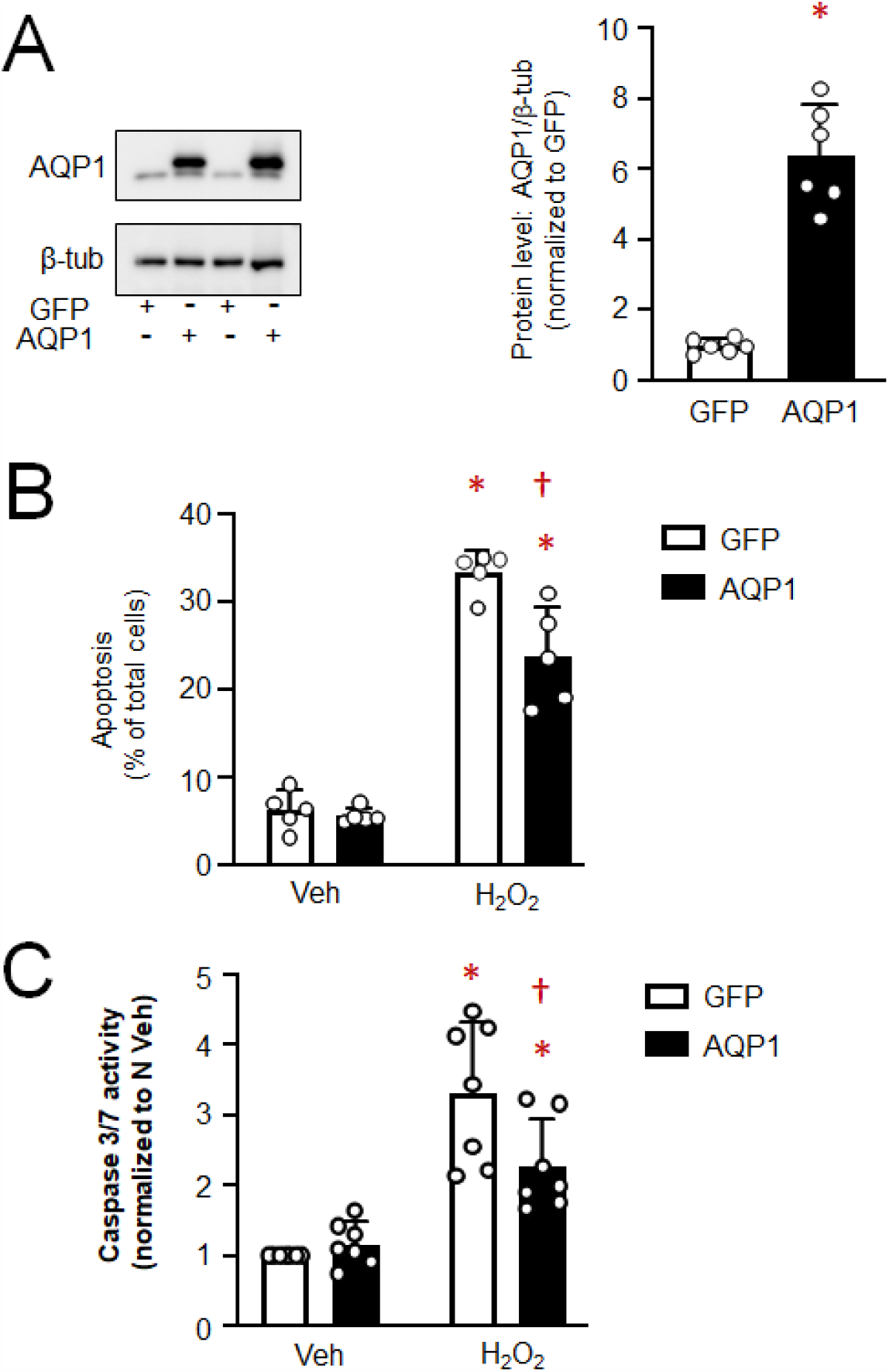
Effect of increasing AQP1 on apoptosis in pulmonary arterial smooth muscle cells (PASMCs) from control rats. **A)** Representative blots show AQP1 protein levels in PASMCs infected with adenovirus containing wild-type AQP1 (AdAQPI) or green fluorescence protein (AdGFP). Bar and scatter plot graphs represent mean ± SD values and individual values (n= 6 rats per group) for AQP1 protein levels. * indicates significant difference (p<0.05) from AdGFP value by t-test. **B and C)** Bar and scatter graphs represent mean ± SD values and individual values for apoptosis measured via **B)** Hoescht staining (n=5 rats per group) and **C)** caspase 3/7 activity assay (n=7 rats per group) in PASMCs from control rats infected with AdGFP or AdAQPI and challenged with vehicle (Veh; PBS) or H_2_O_2_ (500 μM; 24 h). Values for apoptosis are presented as percent of total cells while values for caspase 3/7 activity are normalized to AdGFP Veh cells. * indicates significant difference (p<0.05) from Veh-treated within the same group; † indicates significant difference (p<0.05) from H_2_O_2_ value in cells treated with AdGFP.

### Effects of H_2_O_2_ on BAX/Bcl-2 expression in Normoxic and SuHx PASMCs

To begin to assess the mechanism by which AQP1 protein expression might control caspase 3/7 activity and apoptosis, we measured the levels of Bcl-2 family proteins: Bcl-2 and Bcl-2 like protein 4 (BAX). During apoptosis, levels of the anti-apoptotic protein, Bcl-2, typically decrease, allowing accumulation and/or translocation of pro-apoptotic BAX to mitochondria, where it forms pores through which cytochrome C fragments are released into the cytosol and activate caspase 3. In vehicle-treated cells, Bcl-2 protein levels were significantly higher in PASMCs from SuHx rats compared to controls (**Fig 4A**). Challenge with H_2_O_2_ reduced Bcl-2 protein levels in cells from normoxic animals, although the difference did not reach statistical significance (p=0.1813), whereas Bcl-2 levels in SuHx cells was unaltered by H_2_O_2_. Total BAX protein levels were similar in normoxic and SuHx cells at baseline and did not change with exposure to H_2_O_2_ in either cell type. The ratio of BAX/Bcl-2 expression is a surrogate for apoptotic susceptibility. At baseline (vehicle treatment), the BAX/Bcl-2 ratio was not different in normoxic and SuHx cells. However, because of the trends in Bcl-2 and BAX protein levels, H_2_O_2_ significantly increased the ratio of BAX/Bcl-2 expression in cells from normoxic rats whereas the ratio was significantly lower in H_2_O_2_-challenged cells from SuHx rats (**Fig 4C**).

**Figure 4:**
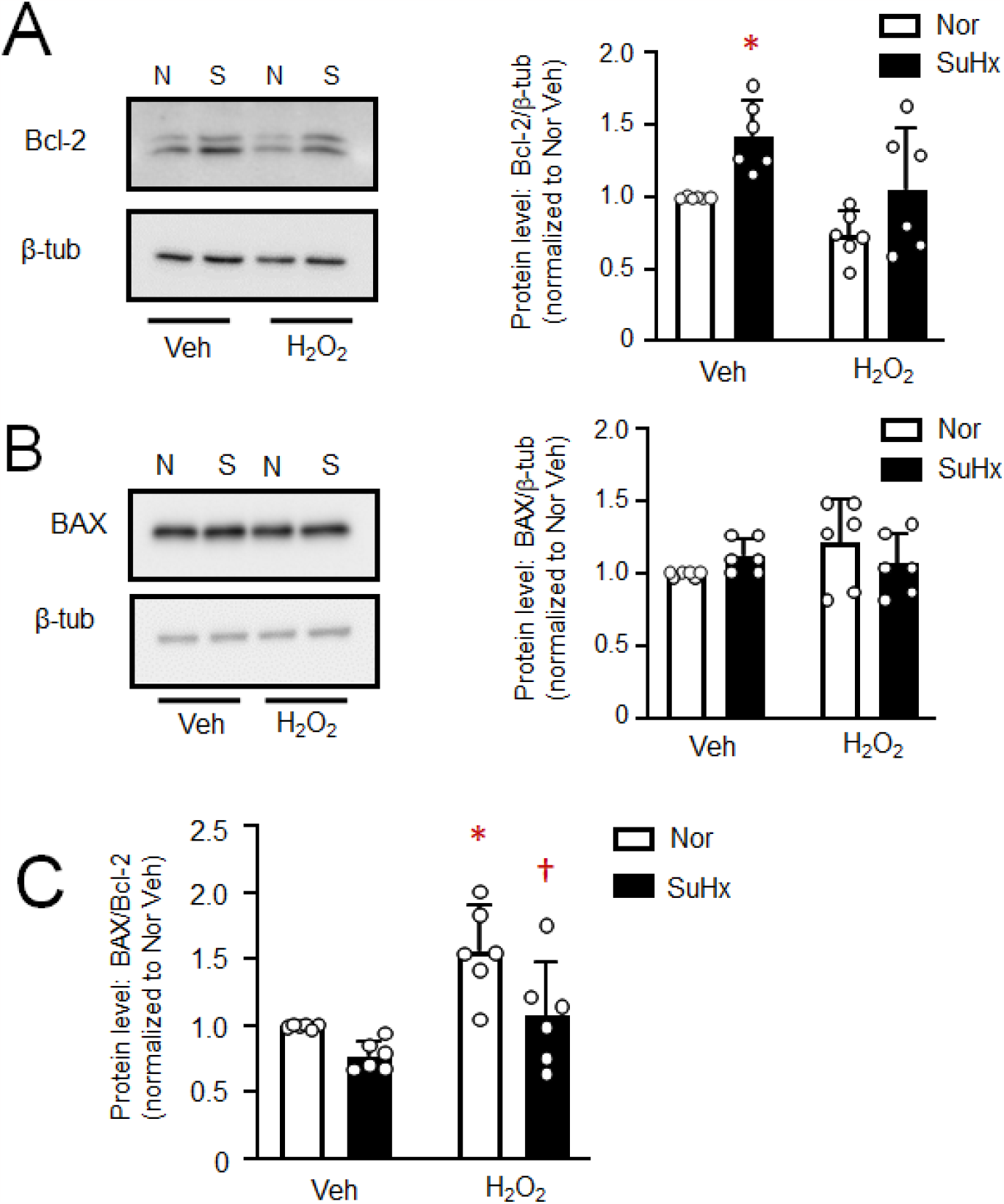
BAX and Bcl-2 levels in pulmonary arterial smooth muscle cells (PASMCs) from normoxic (N) and Su5416+hypoxia (S) rats. **A)** Representative blots show Bcl-2 protein levels in PASMCs from N and S rats treated with vehicle (Veh; PBS) or H_2_O_2_ (500 μM; 24 h). Bar and scatter plot graphs represent mean ± SD values and individual values for Bcl-2 protein levels. * indicates significant difference (p<0.05) from Nor Veh. **B)** Representative blots show BAX protein levels in PASMCs from N and SuHx rats treated with vehicle (Veh; PBS) or H_2_O_2_ (500 μM; 24 h). Bar and scatter plot graphs represent mean ± SD values and individual values for BAX protein levels. **C)** Bar and scatter plot graphs represent mean ± SD values and individual values for the ratio of BAX/Bcl-2 protein levels. For all experiments, n= 6 rats per group * indicates significant difference (p<0.05) from Veh value in the same condition; † indicates significant difference (p<0.05) from H_2_O_2_ value in cells from N rats. Significance assessed by 2-way ANOVA with Holm-Sidak post-hoc test

### Effect of AQP1 depletion on Bcl-2 and BAX protein levels in SuHx PASMCs

Given our finding that the BAX/Bcl-2 ratio following challenge with H_2_O_2_ was significantly reduced in SuHx PASMCs, we next explored whether reducing AQP1 levels in PASMCs from SuHx could restore the effect of H_2_O_2_ on the BAX/Bcl-2 ratio. Depletion of AQP1 in PASMCs from SuHx rats significantly reduced basal Bcl-2 protein levels (**Fig 5A**). Application of H_2_O_2_ reduced Bcl-2 protein levels in SuHx infected with siNT, an effect that was significantly amplified when AQP1 was depleted. Total BAX protein levels remained fairly constant across groups (siNT and siAQP1, with and without H_2_O_2_) (**Fig 5B**). Because of the reductions in Bcl-2, depletion of AQP1 significantly increased the ratio of BAX/Bcl-2 in vehicle-treated SuHx PASMCs at baseline, and significantly enhanced the ability of H_2_O_2_ to increase the ratio of BAX/Bcl-2 protein (**Fig 5C**).

**Figure 5:**
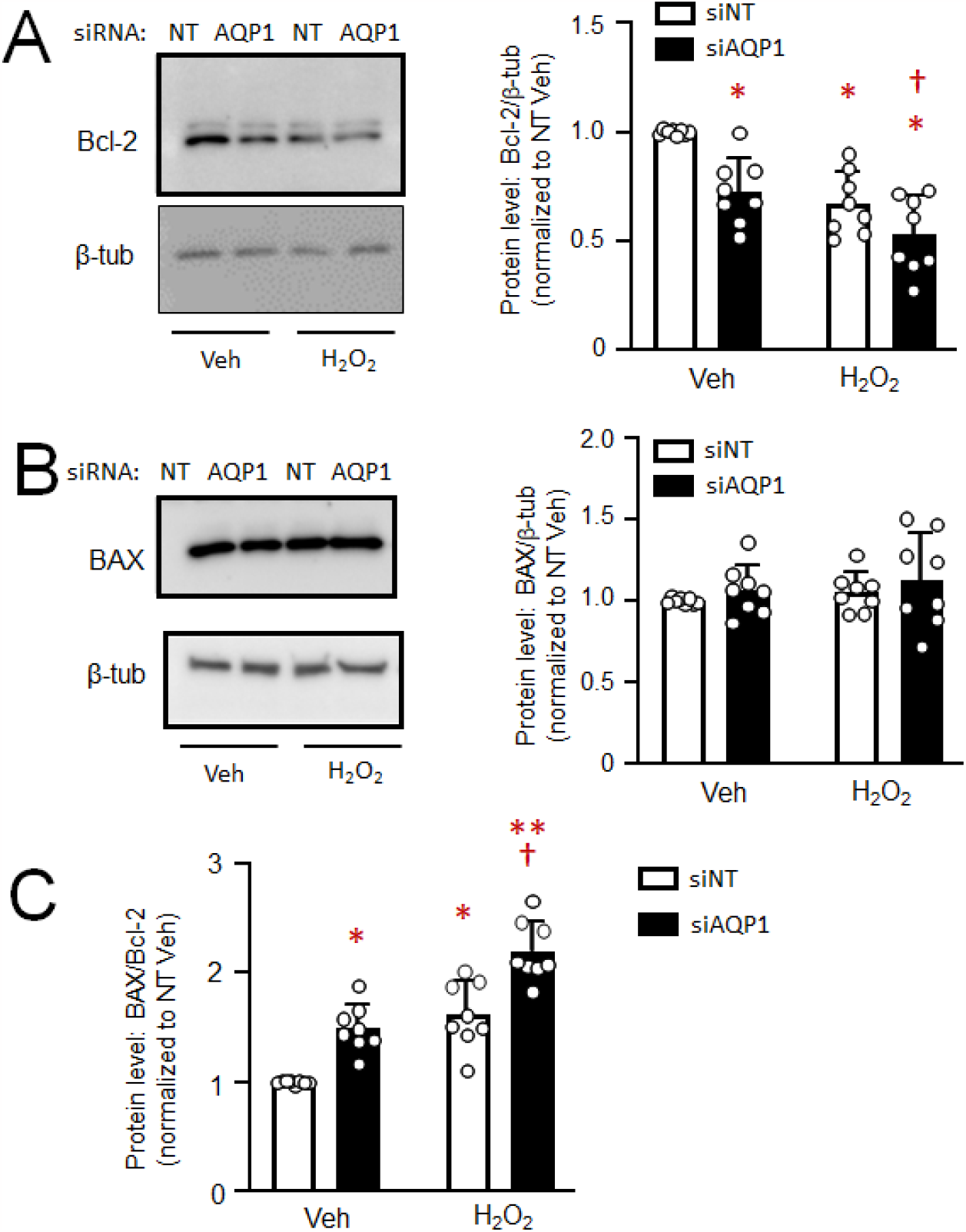
Effect of depleting AQP1 protein on Bcl-2 and BAX protein levels in pulmonary arterial smooth muscle cells (PASMCs) from Su5416+hypoxia (SuHx) rats. **A)** Representative blots show Bcl-2 protein levels in PASMCs from SuHx rats transfected with siRNA targeted to AQP1 (siAQPI) or a non-targeting siRNA (siNT) and treated with vehicle (Veh; PBS) or H_2_O_2_ (500 μM; 24 h). Bar and scatter plot graphs represent mean ± SD values and individual values for Bcl-2 protein levels. * indicates significant difference (p<0.05) from NT Veh; † indicates significant difference (p<0.05) from AQP1 Veh. **B)** Representative blots show BAX protein levels in PASMCs from SuHx rats treated with Veh or H_2_O_2_. Bar and scatter plot graphs represent mean ± SD values and individual values for BAX protein levels. **C)** Bar and scatter plot graphs represent mean ±SD values and individual values for the ratio of BAX/Bcl-2 protein levels. For all experiments, n= 8 rats per group * indicates significant difference (p<0.05) from NT Veh; † indicates significant difference (p<0.05) from AQP1 Veh; ** indicates significant difference (p<0.05) from NT H_2_O_2_. Significance assessed by 2-way ANOVAwith Holm-Sidak post-hoc test.

### Effect of increasing AQP1 protein levels on Bcl-2 and BAX in control PASMCs

In a final set of experiments, we tested whether forced expression of AQP1 was sufficient to alter Bcl-2 and BAX protein expression. In PASMCs from control r ats where we enhanced expression of AQP1, baseline Bcl-2 protein levels were higher than in cells expressing AdGFP, and the H_2_O_2_-induced decrease was attenuated (**Fig 6A**). Increasing AQP1 levels, in the absence or presence of H_2_O_2_, had no effect on total BAX protein levels (**Fig 6B**). The increase in Bcl-2 levels achieved with AdAQP1 was not sufficient to significantly reduced the BAX/Bcl-2 ratio at baseline; however, increasing AQP1 resulted in a significant reduction in the BAX/Bcl-2 ratio in response to H_2_O_2_ compared to H_2_O_2_-challenged cells infected with AdGFP (**Fig 6C**).

**Figure 6:**
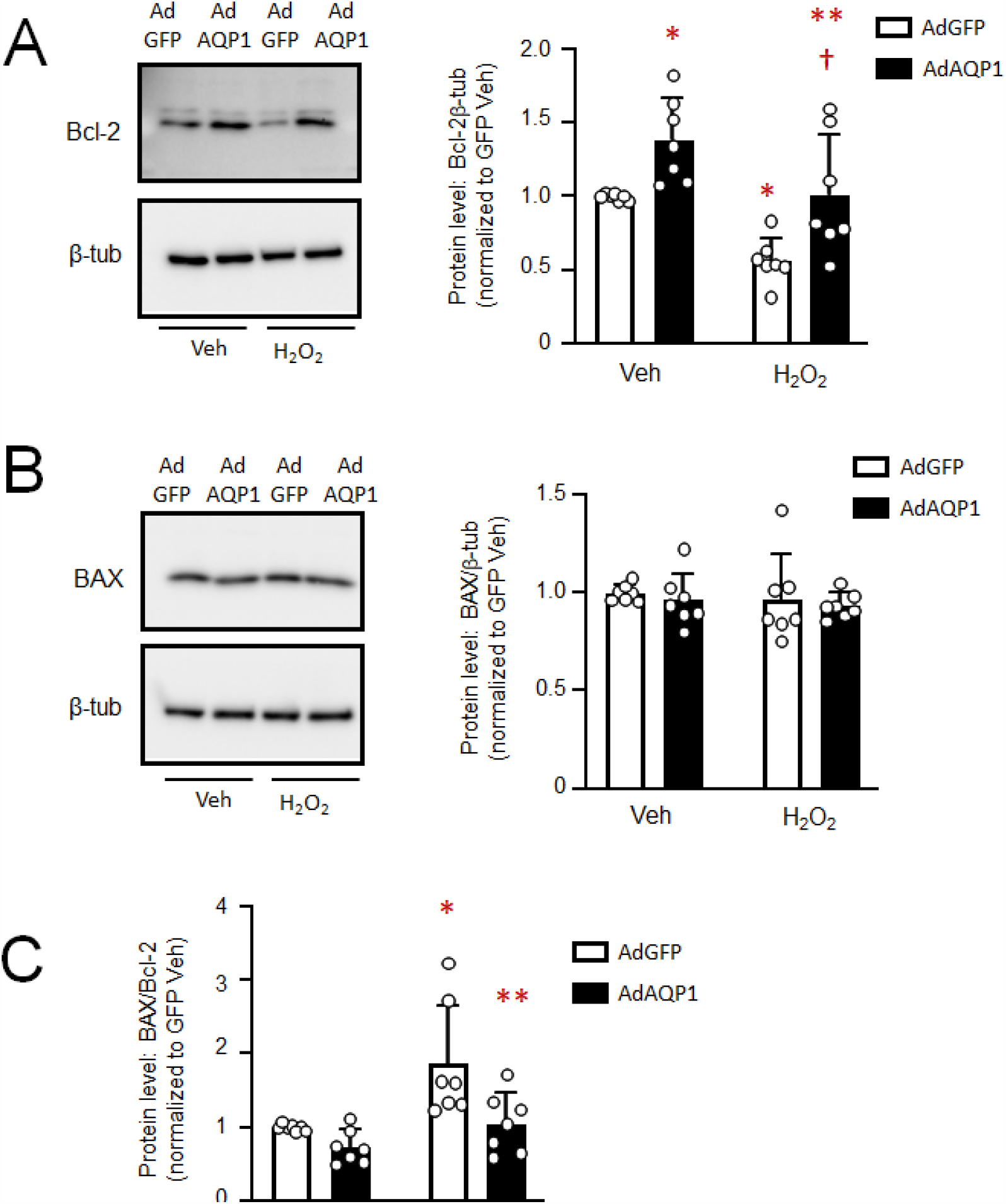
Effect of increasing AQP1 protein levels on Bcl-2 and BAX expression in control pulmonary arterial smooth muscle cells (PASMCs). **A)** Representative blots show Bcl-2 protein levels in PASMCs from control rats infected with adenovirus containing wildtype AQP1 (AdAQPI) or green fluorescent protein (AdGFP) and treated with vehicle (Veh; PBS) or H_2_O_2_ (500 μM; 24 h). Bar and scatter plot graphs represent mean ± SD values and individual values for Bcl-2 protein levels . * indicates significant difference (p<0.05) from GFP Veh; † indicates significant difference (p<0.05) from AQP1 Veh; ** indicates significant difference (p<0.05) from GFP H_2_O_2_. **B)** Representative blots show BAX protein levels in PASMCs from control rats with forced expression of AQP1 or GFP and treated with Veh or H_2_O_2_. Bar and scatter plot graphs represent mean ± SD values and individual values for BAX protein levels. C) Bar and scatter plot graphs represent mean ± SD values and individual values for the ratio of BAX/Bcl-2 protein levels. * indicates significant difference (p<0.05) from GFP Veh; ** indicates significant difference (p<0.05) from GFP H_2_O_2_. Significance assessed by 2-way ANOVA with Holm-Sidak post-hoc test For all experiments, n= 7 rats per group.

## Discussion

In the current study, we tested the role of AQP1 in modulating apoptosis in PASMCs from control cells and in cells from a robust model of PH. We found that elevated levels of AQP1 in PASMCs from pulmonary hypertensive rats were necessary for resistance to apoptosis, and that some level of apoptosis resistance could be conferred simply by increasing expression of AQP1 in PASMCs from control rats. Moreover, in exploring the downstream pathways involved, we found AQP1 levels influence the expression of Bcl-2, with enhanced AQP1 levels corresponding to increased Bcl-2 expression, leading to reductions in the ratio of BAX to Bcl-2 that are typically associated with apoptosis resistance.

It has been widely reported that pulmonary vascular cells from patients with PAH^5,6, 10^ and from pre-clinical models of PH ^5, 31^ are resistant to apoptosis. We confirmed these findings in PASMCs from the SuHx rat model of PH, which recapitulates several of the features of human PAH. Using two complementary methods to assess apoptosis, morphology via Hoechst staining and caspase 3/7 activity, we showed cells from SuHx were resistant to apoptosis induced by H_2_O_2_. Preliminary experiments demonstrated that suppression of apoptosis in these cells was not restricted to a single apoptotic stimulus, as PASMCs from SuHx rats also exhibited reduced apoptosis in response to staurosporine (**Fig S1**).

**Fig S1.**
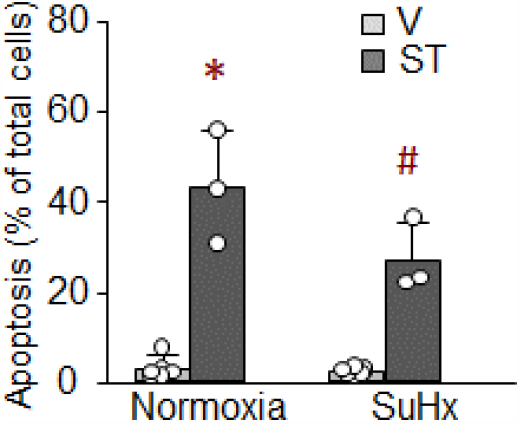
Apoptosis in PASMCs from the SuHx rat model. Cells were stained with Hoechst dye to visualize apoptosis. Bar graphs show mean+/- SD percent apoptotic cells from normoxic and SuHx rats treated with vehicle (V) or staurosporine (ST; 20 nM for 24 h). *p<0.05 from V; # p<0.05 from same treatment in normoxia

An interesting finding from our study was that there was no difference in apoptosis or caspase 3/7 activity at baseline between cells from normoxic and SuHx rats, despite conditions which should produce endogenous pro-apoptotic stimuli, including mitochondrial dysfunction, ER stress, and hyperproliferation^32^,^33^. Lack of increase in baseline apoptosis under these conditions suggests that during PH, PASMCs develop mechanisms to evade activation of executioner caspases. In many types of cancer cells, AQP1 protein levels are elevated and inversely correlated with susceptibility to apoptosis^12^,^28-30^, although in other tumor cells (i.e., mesothelioma) AQP1 and apoptosis were not associated^34^. AQP1 expression was also inversely correlated with apoptosis in aortic endothelial^35^ and lens epithelial^36^ cells. Since SuHx cells also exhibit increased AQP1 expression^18^ (Fig 1) we next tested whether the increased AQP1 levels in SuHx PASMCs was essential for apoptosis resistance. Using siRNA approaches, we showed that reducing the amount of AQP1 in SuHx PASMCs increased apoptosis at baseline and restored the response to H_2_O_2_. These results are consistent with effects reported for hypoxia-induced apoptosis in PASMCs^19^. Although enhanced levels of AQP1 appear to be required to maintain apoptosis resistance in SuHx PASMCs, our results indicate that simply increasing AQP1 levels was not sufficient to fully promote apoptosis resistance. While a reduction in H_2_O_2_-induced apoptosis and caspase 3/7 activity was noted when AQP1 levels were elevated in control PASMCs, the effect was modest, indicating that other factors are also participating in the development of apoptosis resistance during PH. These factors could include increased levels of survivin^31^ or FOXM1^37^, both of which have been associated with apoptosis resistance in PAH. Whether these pathways interact with AQP1 is currently unknown.

Apoptosis can be triggered via intrinsic or extrinsic pathways. In either case, a common component of the pathways is release of fragmented cytochrome C from pores in the mitochondrial membrane formed by the pro-apoptotic Bcl-2 family member, BAX^38,39^. Cytochrome C fragments released into the cytosol activate caspase 3, the executioner caspase, to induce apoptosis. Normally found in the cytosol, BAX activation and translocation to the mitochondria is antagonized by interaction with the pro-apoptotic protein, Bcl-2^38^. Thus, the relative ratio of BAX to Bcl-2 protein can dictate cellular susceptibility to apoptosis, with lower BAX/Bcl-2 ratios associated with reduced apoptosis. In normoxic PASMCs, in response to H_2_O_2_, the increase in the ratio of BAX/Bcl-2 is primarily driven by lower Bcl-2 expression. In PASMCs from SuHx rats, however, there is a trend towards lower BAX/Bcl-2 at baseline, and a reduced increase in the ratio in response to H_2_O_2_. The reduced BAX/Bcl-2 ratio in SuHx cells also appears to be driven primarily by increased Bcl-2 expression, a finding consistent with various tumor cells, where Bcl-2 expression is enhanced, leading to apoptosis resistance^40^. Thus, AQP1-mediated reduction in Bcl-2 expression in SuHx cells provides a mechanism for resisting apoptosis. These results are consistent with findings reported in squamous cell carcinoma, where AQP1 protein levels correlate with reduced Bcl-2 expression^29^.

In contrast to what has often been reported for other cell types^39,41^, we did not observe a change in BAX levels with apoptotic stimulus. While this result was somewhat surprising, some reports have noted that apoptosis is not always accompanied by an increase in BAX^42^. Since we measured total BAX levels, it is possible that observed changes in apoptosis were driven primarily by increased or decreased interaction with Bcl-2, preventing or enhancing, respectively, translocation and formation of mitochondrial pores. Another possibility is that changes in BAX expression occur at an earlier time point than that at which we collected proteins for analysis. Assessing these possibilities will require further experimentation.

While our results demonstrate that AQP1 levels can modulate Bcl-2 expression, the exact mechanism remains unclear. Several reports suggest that β-catenin may play a role. For example, in endothelial cells β-catenin promoted cell survival by upregulating Bcl-2, thus inhibiting BAX activation^43^. Similarly, in myocardial and intestinal epithelial cells, enhancing β-catenin increased Bcl-2 expression^44,45^. We previously reported that AQP1 regulates β-catenin protein levels^21^, through a mechanism involving the C-terminal tail region. Given the plethora of data showing β-catenin regulates Bcl-2 expression, it seems plausible that β-catenin may also be the link between AQP1 and Bcl-2 in PASMCs.

Another possible mechanism by which AQP1 could modulate apoptosis is via water transport. In the earliest stages of apoptosis, one of the most highly conserved events is cell shrinkage due to water loss, or apoptotic volume loss. Consistent with this hypothesis, in granulosa cells, blockade of AQP1 water permeability with mercury prevents cell shrinkage and caspase activation^46^. However, in PASMCs, we observed the opposite effect, with loss of AQP1 promoting apoptosis and activation of caspase 3/7. Whether this reflects the fact that AQP1 levels were reduced, but not completely eliminated, allowing for sufficient water transport to permit cell shrinkage is unknown. Importantly, AQP1 was the only aquaporin identified in granulosa cells, whereas PASMCs also express aquaporins 4 and 7^17^, which may also contribute to water permeability and facilitate apoptotic volume loss to allow apoptosis to proceed when AQP1 abundance is reduced.

AQP1 has also been suggested to facilitate entry of H_2_O_2_ into vascular cells^47^. In this case, depletion of AQP1 could be speculated to prevent apoptosis induced by application of exogenous H_2_O_2_ by reducing H_2_O_2_ levels within the cell. While our experiments cannot entirely rule out this possibility, initial experiments revealed that SuHx cells were also resistant to apoptosis in response to staurosporine, a broad kinase inhibitor (**Fig S1**). Moreover, depletion of AQP1 in SuHx PASMCs significantly increased both basal and staurosporine-induced apoptosis (**Fig S2**). These results would suggest that reducing the speed at which H_2_O_2_ enters the cell is not the main mechanism by which AQP1 confers resistance to apoptosis.

**Fig S2.**
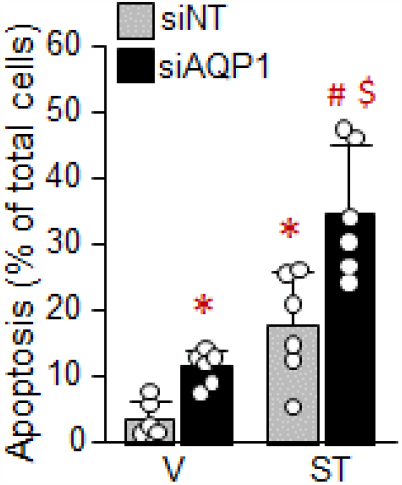
Mean+SD apoptosis (Hoescht stain) in response to staurosporine (ST; 20 nM) or vehicle (V) in PASMCs from SuHx rats treated with SiAQP1 or siNT. *p<0.05 from V: siNT; #p<0.05 from V; siAQPI; $p<0.05 from siNT, ST

In summary, we found that similar to vascular cells from patients with PAH^5,6,10^, PASMCs from the SuHx rat model of PH are resistant to apoptosis via a mechanism involving AQP1-mediated increases in Bcl-2 expression. While necessary for apoptosis resistance observed in these cells, increased AQP1 levels in and of themselves are not sufficient to fully drive apoptosis resistance, suggesting other factors also contribute. These early results provide a mechanism by which AQP1 can regulate PASMC fate and suggest that further investigation could provide additional clues as to whether AQP1-mediated apoptosis resistance contributes to PAH development or progression and whether AQP1 might be a suitable target for therapy.

## Acknowledgments

This work was funded by NIH grants T32 HL007534, F32 HL165766, R01 HL159906, R01 HL126514 and R01 HL073859 and American Heart Association grants 18POST34030262 and 23PRE1022720.

## References

1. Yu Y, Fantozzi I, Remillard CV, Landsberg JW, Kunichika N, Platoshyn O, Tigno DD, Thistlethwaite PA, Rubin LJ and Yuan JX. Enhanced expression of transient receptor potential channels in idiopathic pulmonary arterial hypertension. Proc Natl Acad Sci U S A. 2004;101:13861–6.

2. Morrell NW, Yang X, Upton PD, Jourdan KB, Morgan N, Sheares KK and Trembath RC. Altered growth responses of pulmonary artery smooth muscle cells from patients with primary pulmonary hypertension to transforming growth factor-β1 and bone morphogenetic proteins. Circulation. 2001;104:790–5.

3. Masri FA, Xu W, Comhair SA, Asosingh K, Koo M, Vasanji A, Drazba J, Anand-Apte B and Erzurum SC. Hyperproliferative apoptosis-resistant endothelial cells in idiopathic pulmonary arterial hypertension. Am J Physiol Lung Cell Mol Physiol. 2007;293:L548–54.

4. Jurasz P, Courtman D, Babaie S and Stewart DJ. Role of apoptosis in pulmonary hypertension: from experimental models to clinical trials. Pharmacology & therapeutics. 2010;126:1–8.

5. Meloche J, Le Guen M, Potus F, Vinck J, Ranchoux B, Johnson I, Antigny F, Tremblay E, Breuils-Bonnet S, Perros F, Provencher S and Bonnet S. miR-223 reverses experimental pulmonary arterial hypertension. Am J Physiol Cell Physiol. 2015;309:C363–72.

6. Lampron MC, Vitry G, Nadeau V, Grobs Y, Paradis R, Samson N, Tremblay E, Boucherat O, Meloche J, Bonnet S, Provencher S, Potus F and Paulin R. PIM1 (Moloney Murine Leukemia Provirus Integration Site) Inhibition Decreases the Nonhomologous End-Joining DNA Damage Repair Signaling Pathway in Pulmonary Hypertension. Arterioscler Thromb Vasc Biol. 2020;40:783–801.

7. Stenmark KR, Frid MG, Graham BB and Tuder RM. Dynamic and diverse changes in the functional properties of vascular smooth muscle cells in pulmonary hypertension. Cardiovasc Res. 2018; 114:551 –564.

8. Hua T, Robitaille M, Roberts-Thomson SJ and Monteith GR. The intersection between cysteine proteases, Ca(2+) signalling and cancer cell apoptosis. Biochim Biophys Acta Mol Cell Res. 2023; 1870:119532.

9. Lee YG, Yang N, Chun I, Porazzi P, Carturan A, Paruzzo L, Sauter CT, Guruprasad P, Pajarillo R and Ruella M. Apoptosis: a Janus bifrons in T-cell immunotherapy. J Immunother Cancer. 2023; 11.

10. Dromparis P, Sutendra G and Michelakis ED. The role of mitochondria in pulmonary vascular remodeling. J Mol Med (Berl). 2010;88:1003–10.

11. Hoque MO, Soria JC, Woo J, Lee T, Lee J, Jang SJ, Upadhyay S, Trink B, Monitto C, Desmaze C, Mao L, Sidransky D and Moon C. Aquaporin 1 is overexpressed in lung cancer and stimulates NIH-3T3 cell proliferation and anchorage-independent growth. Am J Pathol. 2006; 168:1345–53.

12. Simone L, Gargano CD, Pisani F, Cibelli A, Mola MG, Frigeri A, Svelto M and Nicchia GP. Aquaporin-1 inhibition reduces metastatic formation in a mouse model of melanoma. J Cell Mol Med. 2018;22:904–912.

13. Zhang Q, Lin L, Li W, Lu G and Li X. MiR-223 inhibitor suppresses proliferation and induces apoptosis of thyroid cancer cells by down-regulating aquaporin-1. J Recept Signal Transduct Res. 2019;39:146–153.

14. Ding T, Zhou Y, Sun K, Jiang W, Li W, Liu X, Tian C, Li Z, Ying G, Fu L, Gu F, Li W and Ma Y. Knockdown a water channel protein, aquaporin-4, induced glioblastoma cell apoptosis. PloS one. 2013;8:e66751.

15. Wu Z, Li S, Liu J, Shi Y, Wang J, Chen D, Luo L, Qian Y, Huang X and Wang H. RNAi-mediated silencing of AQP1 expression inhibited the proliferation, invasion and tumorigenesis of osteosarcoma cells. Cancer biology & therapy. 2015;16:1332–40.

16. Agre P, King LS, Yasui M, Guggino WB, Ottersen OP, Fujiyoshi Y, Engel A and Nielsen S. Aquaporin water channels--from atomic structure to clinical medicine. J Physiol. 2002;542:3–16.

17. Leggett K, Maylor J, Undem C, Lai N, Lu W, Schweitzer KS, King LS, Myers AC, Sylvester JT, Sidhaye VK and Shimoda LA. Hypoxia-induced migration in pulmonary arterial smooth muscle cells requires calcium-dependent upregulation of aquaporin 1. Am J Physiol Lung Cell Mol Physiol. 2012;303(4):L343–53.

18. Yun X, Nm P, Jiang H, Smith Z, Huetsch J, Damarla M, Suresh K and Shimoda LA. Upregulation of aquaporin 1 mediates increased migration and proliferation in pulmonary vascular cells from the rat SU5416/hypoxia model of pulmonary hypertension. Front Physiol. 2021.

19. Liu M, Liu Q, Pei Y, Gong M, Cui X, Pan J, Zhang Y, Liu Y, Liu Y, Yuan X, Zhou H, Chen Y, Sun J, Wang L, Zhang X, Wang R, Li S, Cheng J, Ding Y, Ma T and Yuan Y. Aqp-1 Gene Knockout Attenuates Hypoxic Pulmonary Hypertension of Mice. Arterioscler Thromb Vasc Biol. 2019;39:48–62.

20. Lai N, Lade J, Leggett K, Yun X, Baksh S, Chau E, Crow MT, Sidhaye V, Wang J and Shimoda LA. The aquaporin 1 C-terminal tail is required for migration and growth of pulmonary arterial myocytes. Am J Respir Cell Mol Biol. 2014;50:1010–20.

21. Yun X, Jiang H, Lai N, Wang J and Shimoda LA. Aquaporin 1-mediated changes in pulmonary arterial smooth muscle cell migration and proliferation involve β-catenin. Am J Physiol Lung Cell Mol Physiol. 2017;313:L889–L898.

22. Schuoler C, Haider TJ, Leuenberger C, Vogel J, Ostergaard L, Kwapiszewska G, Kohler M, Gassmann M, Huber LC and Brock M. Aquaporin 1 controls the functional phenotype of pulmonary smooth muscle cells in hypoxia-induced pulmonary hypertension. Basic research in cardiology. 2017;112:30.

23. Edlich F. BCL-2 proteins and apoptosis: Recent insights and unknowns. Biochem Biophys Res Commun. 2018;500:26–34.

24. Moldoveanu T and Czabotar PE. BAX, BAK, and BOK: A Coming of Age for the BCL-2 Family Effector Proteins. Cold Spring Harbor perspectives in biology. 2020;12.

25. Shu C, Shu Y, Gao Y, Chi H and Han J. Inhibitory effect of AQP1 silencing on adhesion and angiogenesis in ectopic endometrial cells of mice with endometriosis through activating the Wnt signaling pathway. Cell cycle. 2019;18:2026–2039.

26. Zhang X, Chen Y, Dong L and Shi B. Effect of selective inhibition of aquaporin 1 on chemotherapy sensitivity of J82 human bladder cancer cells. Oncol Lett. 2018;15:3864–3869.

27. Huetsch JC, Jiang H, Larrain C and Shimoda LA. The Na+/H+ exchanger contributes to increased smooth muscle proliferation and migration in a rat model of pulmonary arterial hypertension. Physiol Rep. 2016;4.

28. Moosavi MS and Elham Y. Aquaporins 1, 3 and 5 in Different Tumors, their Expression, Prognosis Value and Role as New Therapeutic Targets. Pathol Oncol Res. 2020;26:615–625.

29. Lehnerdt GF, Bachmann HS, Adamzik M, Panic A, Koksal E, Weller P, Lang S, Schmid KW, Siffert W and Bankfalvi A. AQP1, AQP5, Bcl-2 and p16 in pharyngeal squamous cell carcinoma. J Laryngol Otol. 2015;129:580–6.

30. Tomita Y, Dorward H, Yool AJ, Smith E, Townsend AR, Price TJ and Hardingham JE. Role of Aquaporin 1 Signalling in Cancer Development and Progression. Int J Mol Sci. 2017;18.

31. McMurtry MS, Archer SL, Altieri DC, Bonnet S, Haromy A, Harry G, Puttagunta L and Michelakis ED. Gene therapy targeting survivin selectively induces pulmonary vascular apoptosis and reverses pulmonary arterial hypertension. J Clin Invest. 2005;115:1479–91.

32. Chen KH, Dasgupta A, Lin J, Potus F, Bonnet S, Iremonger J, Fu J, Mewburn J, Wu D, Dunham-Snary K, Theilmann AL, Jing ZC, Hindmarch C, Ormiston ML, Lawrie A and Archer SL. Epigenetic Dysregulation of the Dynamin-Related Protein 1 Binding Partners MiD49 and MiD51 Increases Mitotic Mitochondrial Fission and Promotes Pulmonary Arterial Hypertension: Mechanistic and Therapeutic Implications. Circulation. 2018;138:287–304.

33. Huetsch J, Yun X, Jiang H and Shimoda L. ER Stress Induced Apoptosis of Smooth Muscle in Pulmonary Hypertension is Regulated by the Sodium-Hydrogen Exchanger. FASEB J. 2019;33:550.1.

34. Klebe S, Griggs K, Cheng Y, Driml J, Henderson DW and Reid G. Blockade of aquaporin 1 inhibits proliferation, motility, and metastatic potential of mesothelioma in vitro but not in an in vivo model. Dis Markers. 2015;2015:286719.

35. Sada K, Nishikawa T, Kukidome D, Yoshinaga T, Kajihara N, Sonoda K, Senokuchi T, Motoshima H, Matsumura T and Araki E. Hyperglycemia Induces Cellular Hypoxia through Production of Mitochondrial ROS Followed by Suppression of Aquaporin-1. PloS one. 2016;11:e0158619.

36. Zheng HH, Xu GX, Guo J, Fu LC and Yao Y. Aquaporin-1 down regulation associated with inhibiting cell viability and inducing apoptosis of human lens epithelial cells. Int J Ophthalmol. 2016;9:15–20.

37. Bourgeois A, Lambert C, Habbout K, Ranchoux B, Paquet-Marceau S, Trinh I, Breuils-Bonnet S, Paradis R, Nadeau V, Paulin R, Provencher S, Bonnet S and Boucherat O. FOXM1 promotes pulmonary artery smooth muscle cell expansion in pulmonary arterial hypertension. J Mol Med (Berl). 2018;96:223–235.

38. Antonsson B, Conti F, Ciavatta A, Montessuit S, Lewis S, Martinou I, Bernasconi L, Bernard A, Mermod JJ, Mazzei G, Maundrell K, Gambale F, Sadoul R and Martinou JC. Inhibition of BAX channel-forming activity by Bcl-2. Science. 1997;277:370–2.

39. Lalier L, Cartron PF, Juin P, Nedelkina S, Manon S, Bechinger B and Vallette FM. BAX activation and mitochondrial insertion during apoptosis. Apoptosis. 2007;12:887–96.

40. Kaloni D, Diepstraten ST, Strasser A and Kelly GL. BCL-2 protein family: attractive targets for cancer therapy. Apoptosis. 2023;28:20–38.

41. Han J, Sabbatini P, Perez D, Rao L, Modha D and White E. The E1B 19K protein blocks apoptosis by interacting with and inhibiting the p53-inducible and death-promoting BAX protein. Genes Dev. 1996;10:461–77.

42. Benito A, Silva M, Grillot D, Nunez G and Fernandez-Luna JL. Apoptosis induced by erythroid differentiation of human leukemia cell lines is inhibited by Bcl-XL. Blood. 1996;87:3837–43.

43. Tajadura V, Hansen MH, Smith J, Charles H, Rickman M, Farrell-Dillon K, Claro V, Warboys C and Ferro A. beta-catenin promotes endothelial survival by regulating eNOS activity and flow-dependent anti-apoptotic gene expression. Cell Death Dis. 2020;11:493.

44. Mezhybovska M, Wikstrom K, Ohd JF and Sjolander A. The inflammatory mediator leukotriene D4 induces beta-catenin signaling and its association with antiapoptotic Bcl-2 in intestinal epithelial cells. J Biol Chem. 2006;281:6776–84.

45. Kaga S, Zhan L, Altaf E and Maulik N. Glycogen synthase kinase-3beta/beta-catenin promotes angiogenic and anti-apoptotic signaling through the induction of VEGF, Bcl-2 and survivin expression in rat ischemic preconditioned myocardium. Journal of molecular and cellular cardiology. 2006;40:138–47.

46. Jablonski EM, Webb AN, McConnell NA, Riley MC and Hughes FM, Jr. Plasma membrane aquaporin activity can affect the rate of apoptosis but is inhibited after apoptotic volume decrease. Am J Physiol Cell Physiol. 2004;286:C975–85.

47. Al Ghouleh I, Frazziano G, Rodriguez AI, Csanyi G, Maniar S, St Croix CM, Kelley EE, Egana LA, Song GJ, Bisello A, Lee YJ and Pagano PJ. Aquaporin 1, Nox1, and Ask1 mediate oxidant-induced smooth muscle cell hypertrophy. Cardiovasc Res. 2013;97:134–42.

